# TRAF6 directs Foxp3 localization and facilitates Treg function through K63-type ubiquitination

**DOI:** 10.1101/316000

**Authors:** Xuhao Ni, Jinhui Tao, Jian Gu, Benjamin V. Park, Zuojia Chen, Stephanie Newman, Hongbin Shen, Xuehao Wang, Bin Li, Bruce R. Blazar, Joseph Barbi, Fan Pan, Ling Lu

**Affiliations:** Liver Transplantation Center, First Affiliated Hospital, Nanjing Medical University, Nanjing, Jiangsu 210029, China; Immunology and Hematopoiesis Division, Department of Oncology, Sidney Kimmel Comprehensive Cancer Center, Johns Hopkins University School of Medicine, Baltimore, Maryland 21287, USA; Key Laboratory of Molecular Virology and Immunology, Unit of Molecular Immunology, Institut Pasteur of Shanghai, Shanghai Institutes for Biological Sciences, Chinese Academy of Sciences, Shanghai 200025, China; Department of Immunology, Roswell Park Cancer Institute, Buffalo, New York 14263, USA; Jiangsu Key Laboratory of Xenotransplantation, Nanjing Medical University, Nanjing, Jiangsu 211166, China; Department of Epidemiology and Biostatistics, School of Public Health, Nanjing Medical University, Nanjing, Jiangsu 211166, China; Department of Pediatrics, Division of Blood and Marrow Transplantation, University of Minnesota, Minneapolis, MN 55455, USA.

**Keywords:** Traf6, Tregs, Foxp3, ubiquitin, K63, tumor

## Abstract

Regulatory T cells (Treg) are crucial mediators of immune control. The characteristic gene expression and suppressive function of Treg depend considerably on the stable expression and activity of the transcription factor Foxp3. While transcriptional regulation of the *Foxp3* gene has been studied in depth, both the expression and function of Foxp3 are also modulated at the protein level. However, the molecular players involved in posttranslational Foxp3 regulation are just beginning to be elucidated. Here we found TRAF6-deficient Tregs were dysfunctional in vivo; mice with Treg-restricted deletion of TRAF6 were resistant to B16 melanomas and displayed enhanced anti-tumor immunity. We further determined that Foxp3 undergoes lysine-63 chain (K63) ubiquitination at lysine 262 mediated by the E3 ligase TRAF6. When deprived of TRAF6 activity or rendered insensitive to K63 ubiquitination, Foxp3 displayed aberrant, perinuclear accumulation, disrupted function. Thus, Foxp3 ubiquitination by TRAF6 ensures proper localization of Foxp3 and facilitates Foxp3’s gene-regulating activity in Tregs. These results implicate TRAF6 as a key posttranslational, Treg-stabilizing force that may be targeted in novel tolerance-breaking therapies.

## Introduction

In a functional immune system, the target and amplitude of immune responses are tightly controlled. Regulatory T cells (Tregs) are among the safeguards that prevent aberrant immune responses including those underlying autoimmunity, inflammatory diseases or allergy (1). This critical function of Tregs is mediated by several suppressive mechanisms including production of anti-inflammatory cytokines (IL-10, TGFβ, IL-35, etc.), expression of co-inhibitory receptor signaling, and the disruption of effector cell growth and metabolism (2). The expansion or enhancement of Tregs may be an effective means to enforce immune tolerance to transplanted grafts or restore homeostasis to patients suffering from autoimmune pathologies (3). Conversely, Treg depletion or inhibition approaches can be exploited to provoke effective anti-tumor immunity in the cancer setting. Therefore, a thorough understanding of the molecules and pathways important for the function of these cells can have broad therapeutic implications.

The suppressive capabilities of Tregs are underpinned by a characteristic gene expression profile defined in large part by the transcription factor Foxp3. This forkhead/wing-helix family member works in concert with multiple co-regulator molecules (Eos, IRF-4, etc.) and a Treg-specific array of epigenetic modifications to shape the transcriptional landscape of Tregs (4-8). This is typified by the stabilized expression of Treg-associated genes (e.g. *Ctla4*, *Il2ra*) and the silencing of genes associated with effector T cell lineages (e.g. *Il2*, *Ifng*). Stable expression of Foxp3 is not only characteristic of Tregs, it is also key for their identity and suppressive function.

While the mechanisms controlling *Foxp3* transcription are important in Treg induction and stability (9), posttranslational regulation controls the stability and activity of the cellular Foxp3 protein pool and hence Tregs.. For example, multiple posttranslational modifications of Foxp3 have significant consequences for Treg function. Acetylation at specific lysine residues improves the stability of Foxp3 expression and enhances the association of Foxp3 with its target genes (10-12). Phosphorylation of Foxp3 at distinct sites has been found to have positive and negative effects on Foxp3 activity and Treg functions (13-14).

We and others have documented how ubiquitination affects the Foxp3 protein pool and the suppressive capacities of Tregs (15-16). Protein ubiquitination regulates diverse biological functions by virtue of both the modified target protein site and nature of the ubiquitin-to-ubiquitin linkages within the added polymer chains. For instance, the addition of ubiquitin monomers linked together at Lysine residue 48 (K48-linkage) often marks a target protein for degradation by the 26S proteasome(17). Other types of ubiquitination, such as those involving K63-linked ubiquitin molecules, can be important for the activation of signaling molecules and protein trafficking events (18-19). Previously, we and others found that Foxp3 was subject to K48-linked polyubiquitination that mediated the transcription factor’s degradation via the proteasome. Invoking or preventing this modification could effectively sabotage (15) or stabilize suppressive function (16), respectively, in Tregs. However, the potentially diverse outcomes of Foxp3 ubiquitination and molecular pathways involved have been incompletely elucidated.

Here we report a distinct pathway of K63-type Foxp3 ubiquitination capable of promoting Foxp3 nuclear localization, regulatory activity and Treg stability and function, mediated by the TNF receptor associated factor protein family member, TRAF6. Mice lacking TRAF6 specifically in Foxp3+ Tregs displayed compromised immune tolerance and robust immune activation at baseline compared to wild type controls(17-23). But the mechanism of the regulation of TRAF6 to Foxp3 was not determined. We found such mice were resistant to aggressive melanomas accompanied by aberrant extranuclear accumulation of Foxp3 and robust anti-tumor immune responses. Furthermore, we show that the zinc finger and leucine zipper domains of Foxp3 and lysine residue 262 play a critical role in interacting with Traf6 and the process of K63- ubiquitination. Taken together, these results demonstrate a hitherto unknown, posttranslational mechanism controlling both the regulatory activity of Foxp3 and Treg suppressive potency and identify TRAF6 as a potential target for focused, tolerance-breaking immunotherapies.

## Results

### Traf6 is critical for enforcement of immune homeostasis by Tregs

TRAF6 plays a potentially significant role in Treg biology. In line with this notion, high levels of TRAF6 transcript were found in naïve CD4+ T cells differentiating into Tregs in vitro (induced or iTregs), but not those committing to other T helper lineages (**Fig.Fig. 1A**). Our previous study determined that miRNA-146b-5p increased TRAF6 expression in human tTregs (24). We further found that nTregs freshly isolated from human peripheral blood expressed TRAF6 mRNA to a greater degree than their non-Treg CD4+ counterparts (**Fig.Fig. 1B**). This preferential expression of TRAF6 by multiple Treg subsets further implicates the E3 ligase as a key factor in the development and biology of these important suppressor cells.

**Fig. 1.**
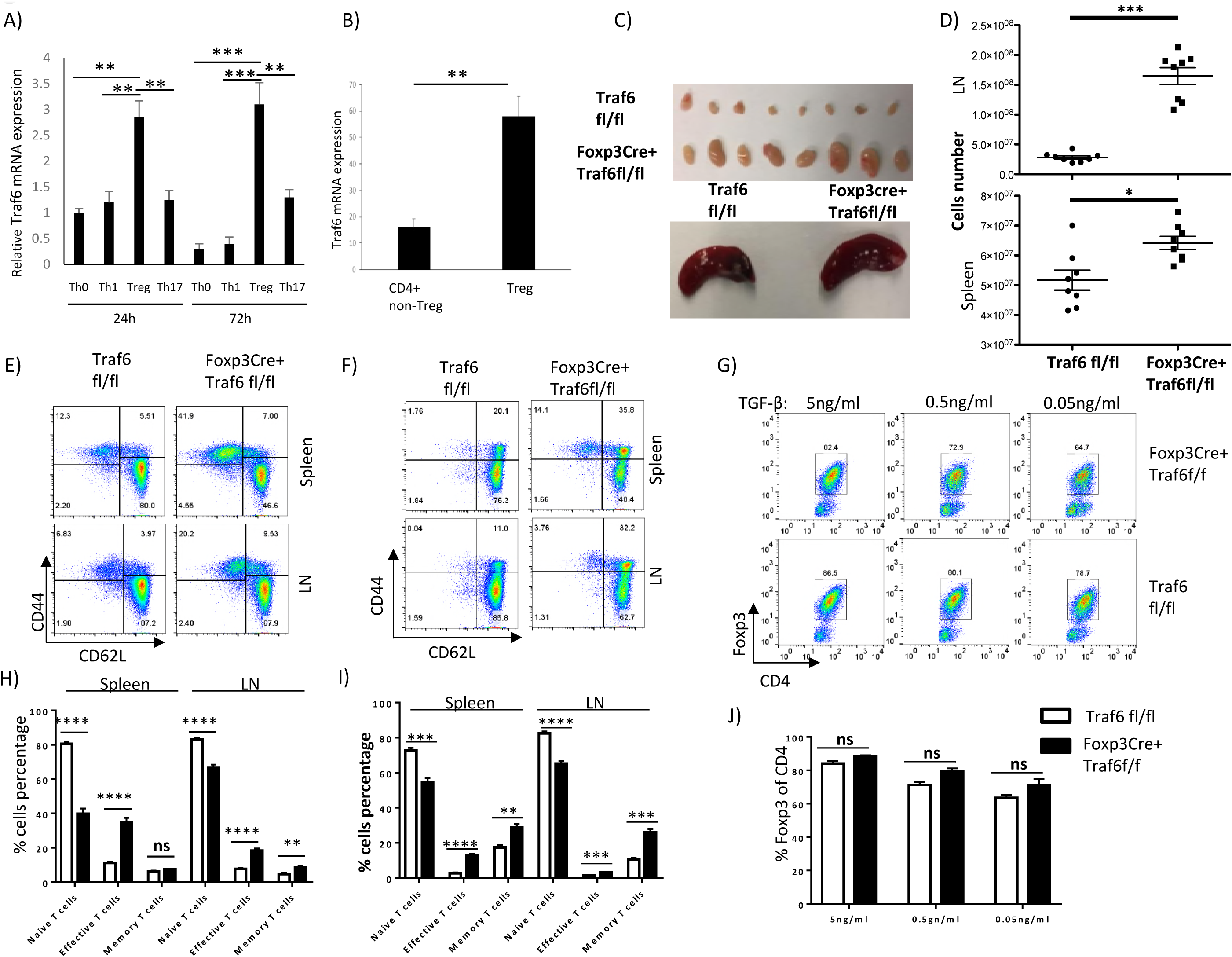
TRAF6 is highly expressed by Treg subsets and plays an important role in maintaining immune homeostasis. (A) TRAF6 expression in differentiating CD4+ T cells. Naïve CD4+ T cells were obtained from wild type C57BL/6 mice by FACS and activated with anti-CD3/CD28 (1μg, and 2μg/ml) for the indicated times in the presence of distinct Thelper lineage-directing cytokines or under neutral activation conditions (Th0). After total RNA extraction and cDNA conversion, RT-PCR determined of TRAF6 mRNA in differentiating Th0, Th1, Th17 and iTregs was determined. (B) TRAF6 mRNA expression by human Tregs and non-Treg CD4+ T cell. Human Tregs (CD3^+^/CD4^+^/CD8^−^/CD25^HIGH^/CD127^low^/CD39^+^) and non-Treg CD4+ T cells were obtained from the peripheral blood of healthy donors by FACS after Ficoll-Paque PLUS gradient centrifugation and magnetic bead enrichment of CD4+ T cells. TRAF6 mRNA was measured by qRT-PCR. (C, D) Evidence of lymphoproliferative disease in Traf6^fl/fl^Foxp3Cre^+^ mice. (C) Spleens and lymph nodes were recovered from Traf6^fl/fl^Foxp3Cre^+^ mice and Traf6^fl/fl^ littermates at 8 weeks of age. (D) The cellularity of the lymphoid tissues of Traf6^fl/fl^Foxp3Cre^+^ mice and their Traf6^fl/fl^ littermates was determined (8 mice/group). (E, H) Effect of Treg-specific TRAF6-deficiency on baseline T cell activation. The frequencies of the effective cells (CD44^high^/CD62L^low^), the memory cells (CD44^high^/CD62L^high^) and the naïve cells (CD44^low^/CD62L^high^) in the CD4+ T cell compartments of Traf6^fl/fl^ and Traf6^fl/fl^Foxp3Cre^+^ mice were determined by flow cytometry (5 mice/group). (F, I) Effect of Treg-specific TRAF6-deficiency on baseline T cell activation. The frequencies of the effective cells (CD44^high^/CD62L^low^), the memory cells (CD44^high^/CD62L^high^) and the naïve cells (CD44^low^/CD62L^high^) in the CD8+ T cell compartments of Traf6^fl/fl^ and Traf6^fl/fl^Foxp3Cre^+^ mice were determined by flow cytometry (5 mice/group). (G, J) Impact of TRAF6-expression on *in vitro* Treg differentiation. As in A, naïve CD4+ T cells were isolated from Traf6^fl/fl^Foxp3Cre^+^ and Traf6^fl/fl^ mice and differentiated into iTregs. Conditions of sub-optimal TGFβ concentrations (0.5ng/ml, 0.05ng/ml) were tested as well and intracellular Foxp3 was measured after 4 days, three independent experiments performed. Panels B, D, H, I and J represent the mean results +/-SEM. \**P*<0.05; \*\**P*<0.01; \*\*\**P*<0.001; \*\*\*\**P*<0.0001; ns, no significance, unpaired t test. A, C, E-G are representative of 3 experiments.

To further investigate the importance of TRAF6 for Treg identity and function, we explored the consequences of TRAF6-deficiency in vivo. Previously, global TRAF6 knockout was found to precipitate multi-organ autoimmunity (23). Similarly, T cell-specific deletion of TRAF6 results in lymphoproliferative disease and systemic immune activation tied to T cells resistant to Treg suppression (25). We set out to determine how much of this immune dysregulation stems from the absence of TRAF6 activity in Tregs by observing the consequences of Foxp3-driven TRAF6 deletion. Here, mice possessing a TRAF6 gene flanked with LoxP sites were crossed to mice expressing the cre-recombinase under the control of the Foxp3 promoter. Resulting Treg-specific knockouts (Traf6^fl/fl^Foxp3Cre^+^) and their wild type littermates (Traf6^fl/fl^Foxp3Cre^−^, “WT”) were monitored for indications of disrupted immune control. Indeed, Traf6^fl/fl^Foxp3Cre^+^ mice displayed a signs of lymphoproliferative disease in lymph nodes and spleens that were noticeably enlarged relative to wild type littermates (**Fig.Fig. 1C**). Increased cellularity was also noted in these lymphoid tissues in the absence of Treg-derived TRAF6 (**Fig.Fig. 1D**). Flow cytometry analysis of lymphocyte surface markers revealed that both the CD4+ and CD8+ T cell compartments of Traf6^fl/fl^Foxp3Cre^+^ mice harbored greater proportion of cells displaying an activated surface marker profile (CD44high/CD62Llow) and fewer resting/naïve (CD44low/CD62Lhigh) cells, indicative of enhanced baseline immune activation (**Fig. 1E, 1H, 1F, 1I**). Furthermore, the frequencies of cells producing proinflammatory cytokines (IFNγ, IL-17) were noticeably increased in the lymph nodes and spleens of Traf6^fl/fl^Foxp3Cre^+^ mice relative to WT controls at baseline (**Fig. S1A-1B**). Commensurate with these indications of poorly enforced immune tolerance and a propensity towards T cell activation, and in line with another recent study (22), Traf6^fl/fl^Foxp3Cre^+^ mice display stunted weight gain with age compared to their wild type littermates (data not shown).

These observations clearly illustrate the importance of TRAF6 in the broad maintenance of immune tolerance by Tregs. As stable and robust Foxp3 expression is central to the suppressive phenotype of Tregs, a positive role for TRAF6 in the activation or maintenance of this factor could explain the effects of knocking out this enzyme. Indeed, prior work has implicated TRAF6 as important for thymic Treg generation and Foxp3 expression in peripheral lymphoid tissues (23) and reduced expression of Foxp3 in TRAF6-deficeit Tregs (22). These results suggest that TRAF6 expression in the Treg compartment is necessary for an optimal pool of these suppressor cells.

We next explored the effects of Treg-specific TRAF6 knockout on Treg differentiation. Prior work has shown that T cell-restricted TRAF6 deficiency does not impact in vitro, TGFβ-driven induction of Foxp3 expression (22). Also, despite displaying stunted thymic Treg development, mice globally deficient in TRAF6 yield T cells more inclined to Foxp3 induction *in vitro* (23). We found that in vitro activation of naïve CD4+ T cells from wild type and Traf6^fl/fl^Foxp3Cre^+^ mice in the presence of TGFβ and IL-2 resulted in similar Foxp3-induction, even with sub-optimal concentrations of TGFβ (**Fig. 1G and 1J**). However, in other experiments, addition of the proinflammatory, Th17-inducing cytokine, IL-6 to strongly Treg-inducing media (5 ng/ml TGFβ) resulted in disrupted iTreg skewing evidenced by low levels of Foxp3 induction. This was seen to an even greater degree when TRAF6 was deleted in the newly induced Tregs (**Fig. S2A and 2B**). Interestingly, when fully differentiated iTregs, which can be prone to unstable Foxp3 expression, were treated with proinflammatory cytokines FoxP3 expression decreased more readily in the absence of TRAF6 expression. This was particularly the case of IL-6 exposure while TNFα and IL-1β treatment reduced Foxp3 expression in both groups. Treatment with the TLR ligand LPS also disproportionately reduced Foxp3 levels in Traf6^fl/fl^Foxp3Cre^+^ derived iTregs (**Fig. S2C-E).** These findings suggest that TRAF6 expression in Tregs may stabilize or bolster expression of Foxp3 in multiple Treg subsets.

### Expression of TRAF6 is required for in vivo Treg suppression

We next investigated the role of TRAF6 activity on the suppressive function of Tregs. To this end we isolated Tregs from wild type (Foxp3cre^+^) and Traf6^fl/fl^Foxp3Cre^+^ mice and compared their suppressive potency using an in vitro suppression assay. The ability of these Tregs to suppress the proliferation of naïve CD4+ “responder” T cells was determined by measuring CFSE-dilution by flow cytometry. We found that TRAF6-deficient Tregs were as functional as wild type Tregs in vitro (**Fig. S3A-B**) in agreement with prior studies of Tregs isolated from Traf6^fl/fl^CD4Cre^+^ mice (25). In contrast, an in vivo assay of Treg function revealed that without TRAF6, Tregs failed to restraint responder T cell expansion. Here, CD45.2+ Tregs from either Traf6^fl/fl^Foxp3Cre^+^ and wild type mice were purified and mixed with normal naïve responder CD4+ T cells (CD45.1+) at a 1:5 ratio prior to injection into lymphopenic Rag2-/- recipient mice, which also agree with prior studies (22). After 7 days, the relative frequencies and numbers of responder and Treg cells in spleen were observed by flow cytometry. Responder cells injected with Traf6^fl/fl^Foxp3Cre^+^−derived Tregs were much more plentiful than those co-transferred with wild type Tregs, and knockout Tregs were scarcer than their wild type counterparts (**Fig. 2A**). These results suggest that TRAF6 is necessary for Treg suppressive function in vivo. Interestingly, this dysfunction seen in the absence of TRAF6 was not accompanied by disrupted expression of Treg-associated factors. For instance, levels of GITR and CTLA-4 were actually elevated in the Foxp3+ cells of Traf6^fl/fl^Foxp3Cre^+^ mice relative to WT controls. These increases likely reflected a state of hyper-activation akin to that seen across the broader T cell compartment. Staining patterns of CD44 and CD62L on Traf6^fl/fl^Foxp3Cre^+^-derived cells were consistent with a more activated state (**Fig. S3C**), but not necessarily functional Tregs. This could fit in with going from biological outcomes to discerning mechanisms.

**Fig. 2.**
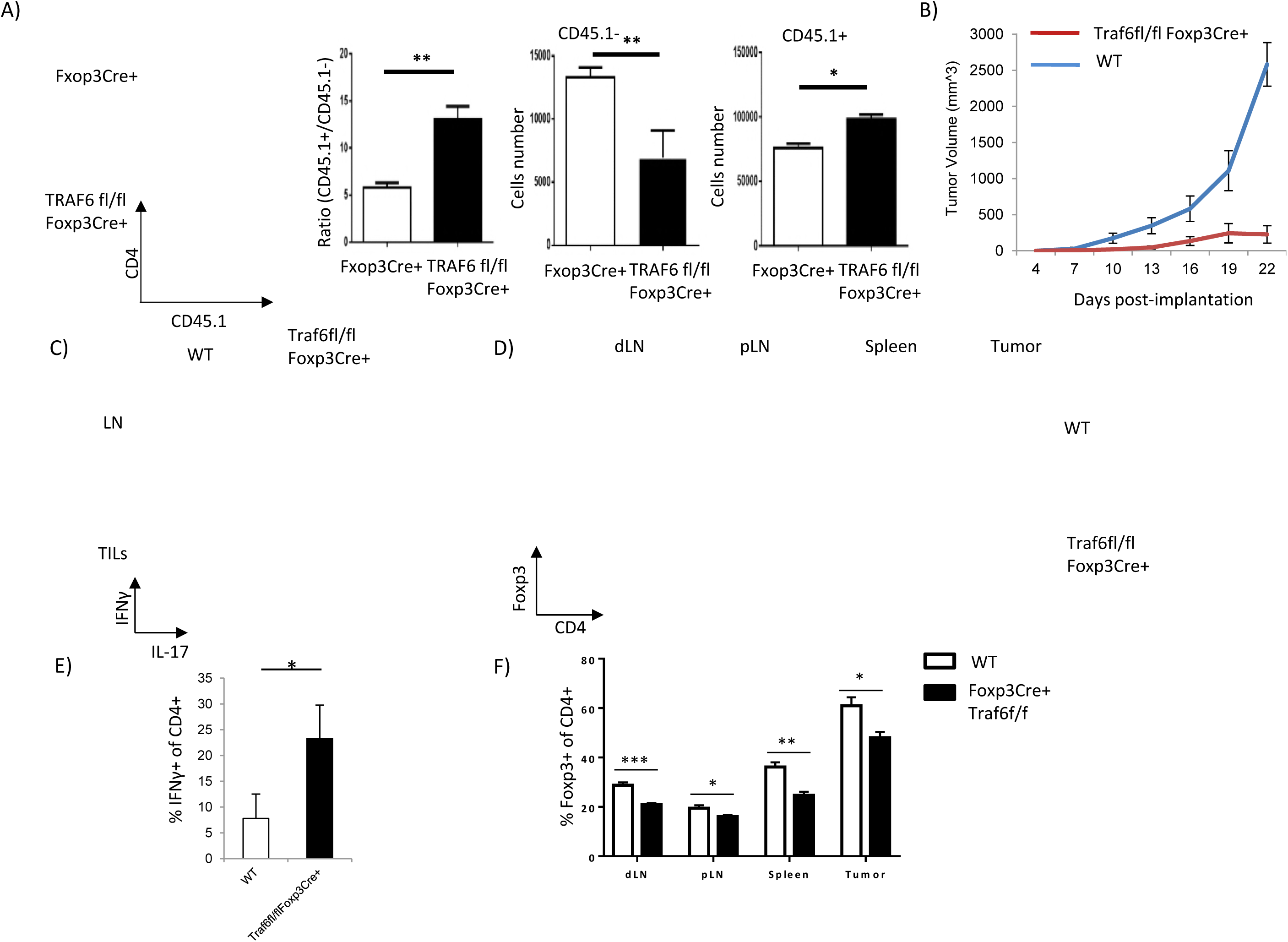
Expression of TRAF6 by Tregs is required for *in vivo* function and suppression of anti-tumor immunity. (A) In vivo suppressive function of WT and Traf6^fl/fl^Foxp3Cre^+^ Tregs. Tregs from the indicated mice (CD45.2+) were isolated by FACS, as were naïve CD4+ responder T cells from congenically distinct (CD45.1+) donor mice. Tregs and Tresponders were mixed at a 1:5 (2×10^5^:10×10^5^) ratio before injection into Rag2-/- mice. 7 days later, spleens were harvested and the relative frequencies and numbers of each transferred cell populations were determined by flow cytometry. (B) Implanted B16 melanoma growth in Traf6^fl/fl^Foxp3Cre^+^ and WT mice. 1×10^5^ B16 cells were injected s.c. into the shaved flanks of the indicated mice (n= 4-7/group). Tumor volumes were monitored every 3 days. (C, E) Proinflammatory cytokine production in tumor-bearing mice with and without Treg-specific TRAF6 expression. Cell suspensions of the tumor-draining lymph node and tumor-infiltrating leukocytes (TILs) were recovered after 21 days of tumor growth. Ex vivo stimulation with PMA and ionomycin in the presence of Golgistop for 5 hour preceded intracellular staining for IFNγ and IL-17 and flow cytometry analysis. (D, F) Foxp3 expression by CD4+ T cells in tumor bearing Traf6^fl/fl^Foxp3Cre^+^ and WT mice. The frequency of Foxp3+ cells in the CD4+ cells of lymph nodes, spleens, and TILs were determined by flow cytometry. \**P*<0.05; \*\**P*<0.01; \*\*\**P*<0.001, unpaired t test. Shown are the representative findings from at least 3 independent experiments (Panels A, C, and D). Error bars in A (right panels), B, E and F denote +/- SEM.

### TRAF6 deficiency enhances anti-tumor immunity and slows the progression of B16 melanomas

We next explored how TRAF6 modulation could affect Treg-enforced immune suppression in the cancer setting. To this end, we challenged Traf6^fl/fl^Foxp3Cre^+^ mice and their TRAF6-competent littermates with aggressive, poorly immunogenic B16 melanomas. After subcutaneous (s.c.) injection, we measured the growth of the implanted tumors. Confirming the inability of TRAF6-deficient Tregs to enforce tumor-associated immune tolerance, Traf6^fl/fl^Foxp3Cre^+^ mice failed to support the growth of implanted B16 melanoma cells. The tumors of wild type mice, on the other hand, readily developed and grew progressively (**Fig. 2B**). In keeping with the notion that TRAF6 is necessary for Treg-mediated immune restrain, the severely stunted tumor progression in Traf6^fl/fl^Foxp3Cre^+^ mice coincided with a markedly enhanced anti-tumor response evidenced by a heightened production of the proinflammatory cytokines IFNγ and IL-17 by leukocytes infiltrating the tumor and tumor-draining lymph node (**Fig. 2C and 2E**). Additionally, deficiencies in Treg TRAF6 expression lead to reduced Foxp3+ T cell frequencies within the CD4+ T cells infiltrating several lymphoid and tumor tissues (**Fig. 2D and 2F**). These findings clearly illustrate the important role played by TRAF6 in Treg-mediated immune control and tumor-enforced suppression of anti-cancer immunity. These findings suggest that potential therapies targeting TRAF6 will be highly effective at breaking tolerance and bolstering the anti-tumor immunity by undermining Treg function.

### Traf6 interacts with Foxp3 and mediates K63 polyubiquitination

Posttranslational modifications can impact the expression and function of Foxp3 in Tregs, and several examples recently have been described (26, 27). Previously, we found that Foxp3 was subject to K48- linked polyubiquitination at lysines 227, 250, 263 and 268, resulting in FoxP3 degradation (15). Since the tagging of target proteins with K48-linked polyubiquitin chains by E3 ligases generally leads to degradation, modification of proteins with chains of ubiquitins inter-linked at other lysine residues (K63 for instance) can yield non-proteolytic outcomes. Such modifications are typically associated with altered intracellular trafficking or activity of target proteins, known to be the case for a number of factors with demonstrated importance to leukocyte function and immune activation (28). Prior to our finding that TRAF6 is apparently capable of promoting Foxp3 ubiquitination, the enzyme was widely known as an important facilitator of K63 ubiquitination (29-31). Potential K63-type modification of Foxp3 and its consequences for Treg function have yet to be studied. To better understand the mechanisms underlying TRAF6 ubiquitin-mediated regulation of Foxp3 function in Tregs, we therefore explored if TRAF6 modifies Foxp3 in this way, and we set out to determine whether this is among the largely unknown mechanisms governing Foxp3 trafficking, nuclear localization, and subsequent gene-regulating activity. To this end, we co-expressed TRAF6 and Foxp3 in cell lines that also expressed a particular variant of Flag-labeled ubiquitin – one in which all lysine residues had been lost (mutated to ubiquitin-resistant arginine residues) save for K63 (“Flag-Ub-K63”). Indeed, robust ubiquitination of Foxp3 was observed only when TRAF6 was expressed. Furthermore, nullifying the E3 ligase activity of TRAF6 through mutation of its RING domain (C70A) mostly ablated the ability of this enzyme to induce K63-type ubiquitination of Foxp3 in cell lines (**Fig. 3A**). These findings implicate TRAF6 as an E3 ligase responsible for K63-linked ubiquitination of Foxp3.

**Fig. 3.**
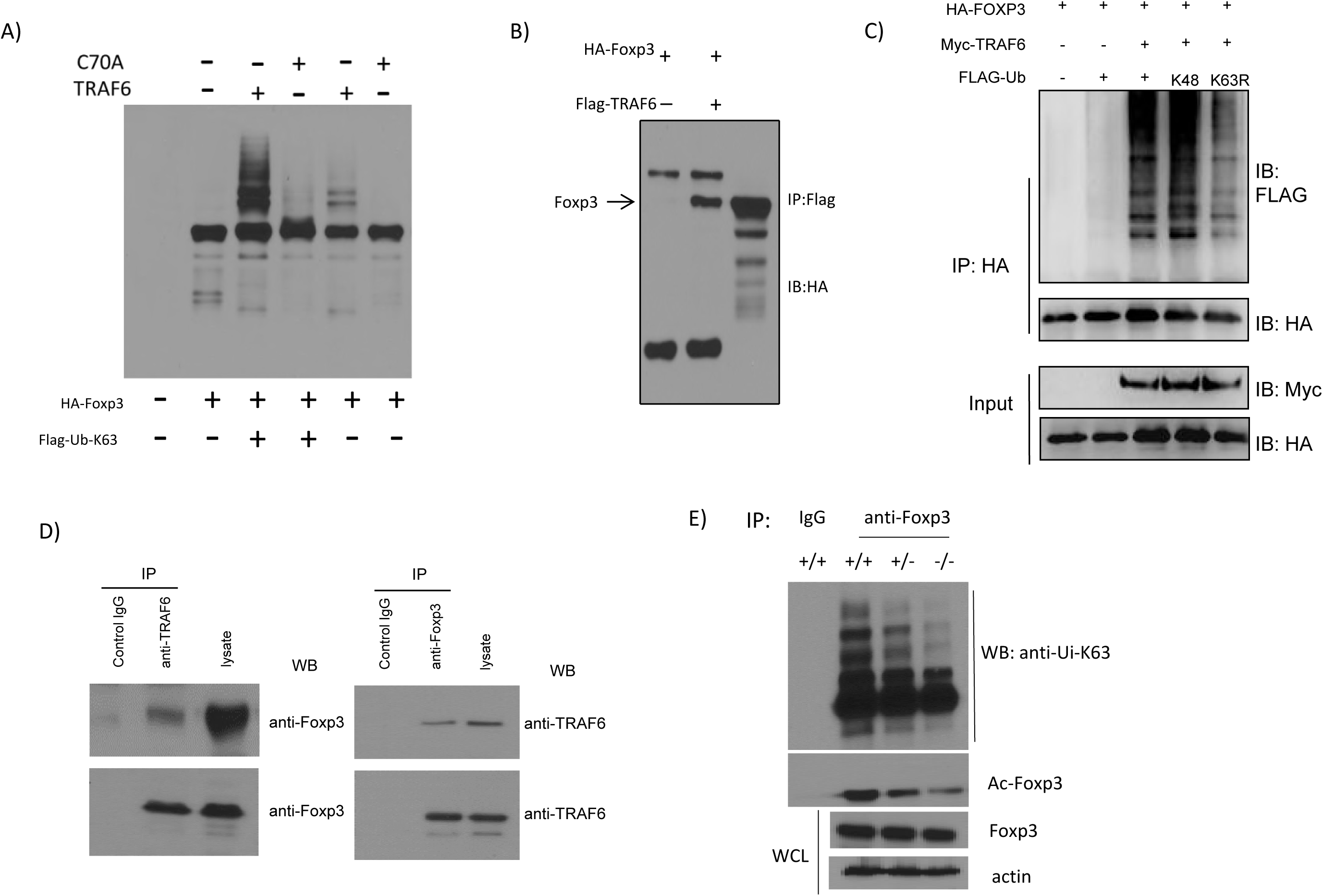
TRAF6 physically interacts with Foxp3 and induces K63-linked polyubiquitination of the transcription factor. (A) Assessment of K63-type ubiquitination of Foxp3 upon expression of wild type and catalytically deficient TRAF6. 293T cells were transfected with the given combinations of vectors encoding wild type TRAF6, the C70A mutant, HA-Foxp3 and Flag-labeled ubiquitin molecules possessing a single lysine residue (K63). (B) Co-immunoprecipitation (co-IP) of Foxp3 with TRAF6 and its facilitation of K63-linked ubiquitination. 293T cells expressing constructs encoding the indicated genes were lysed and incubated with Flag beads to pull down Foxp3, which was detected by immunoblotting. (C) Cell lines expressing the indicated combination of HA-Foxp3, Myc-TRAF6, and Flag-tagged ubiquitin – either wild type Ubi or Ubi-K48 only or Ubi-K63-to-R mutant. (D) Endogenous, reciprocal co-IP of Foxp3 and TRAF6 in murine iTreg lysate. Naïve CD62L^high^/CD25-/CD4+ T cells were obtained by FACS and activated with anti-CD3/CD28 antibodies in the presence of IL-2 and TGFβ for four days before lysis. Antibodies against TRAF6 (left) or Foxp3 (right) were used to pull down target proteins and their interaction partners that were visualized by immunoblotting. (E) Degree of K63 ubiquitination in the cellular Foxp3 pools of Traf6^fl/fl^Foxp3Cre^+^, Traf6^fl/wt^Foxp3Cre^+^ mice and Traf6^wt/wt^Foxp3Cre^+^mice. CD4+/YFP+ cells from each strain were lysed and Foxp3 was immunoprecipitated and levels of acetylated protein was determined by Westernblot analysis as were levels of total Foxp3 and actin. Shown are the representative findings of at least three independent experiments.

To further explore the potential ligase-target relationship between TRAF6 and Foxp3, we tested whether these factors physically interact. Here we performed co-immunoprecipitation (co-IP) experiments with the lysates of cell lines transfected with expression vectors for HA-Foxp3 and FLAG-TRAF6. Upon immunoprecipitation of TRAF6, we detected associated Foxp3 protein among the material recovered from lysate (**Fig. 3B**), suggesting that TRAF6 indeed interacts with Foxp3.

Co-expression of HA-Foxp3 and Myc-TRAF6 constructs along with those encoding distinct FLAG-tagged ubiquitin variants allowed us to confirm the importance of the TRAF6 for this type of Foxp3 modification. Considerable ubiquitination of Foxp3 was seen in cells triple-transfected with expression constructs encoding Foxp3, TRAF6, and wild type ubiquitin (**Fig. 3C**). Those lacking ectopic TRAF6 expression, however, showed only minor levels of Foxp3 polyubiquitination. Interestingly, expression of ubiquitin molecules possessing mutated K48 residues (K48R), which are incapable of supporting K48 polyubiquitination, still resulted in robust Foxp3 ubiquitination in this system. Constructs encoding ubiquitins having a lysine-to-arginine (K-to-R) mutation at residue K63 (K63R), on the other hand, led to a marked reduction in Foxp3 ubiquitination as detected by anti-FLAG immunoblotting after Foxp3 pull-down. These findings demonstrate that TRAF6 plays an important role in facilitating Foxp3 ubiquitination, particularly in K63-type modifications, and they further suggest that such posttranslational modification may be prevalent under baseline conditions. This notion was confirmed in primary murine iTregs using a reciprocal endogenous TRAF6 and FoxP3 co-IP approach (**Fig. 3D**). Importantly, we also found that the dysfunctional TRAF6-deficient Tregs displayed a relative dearth of K63-ubiquitinated Foxp3 species compared to normal Tregs (**Fig. 3E**), further implicating this modification of Foxp3 by TRAF6 as critical for optimal Treg function.

### The zinc finger and leucine zipper domains of Foxp3 and lysine residue 262 are necessary for interaction with Traf6 and ubiquitination

We then set out to further characterize the interaction between Foxp3 and TRAF6. To identify the TRAF6-interacting domain of Foxp3, we generated Foxp3 deletion mutants each lacking one or several of the recognized functional domains of the transcription factor (**Fig. S4A-C**). The ability of full-length Foxp3 and these deletion variants to interact with TRAF6 was determined by co-IP approaches. Molecules that contained intact Zinc-finger and Leucine zipper domains (full length Foxp3, N1, N2, and C3) readily pulled down TRAF6. Meanwhile those lacking these domains (N3, C1, and C2) failed to do so (**Fig. S4D**). These revealed that the Zinc-finger and Leucine zipper domain of Foxp3 were indispensable for the interaction of TRAF6 with Foxp3.

In order to identify potential target sites in the Foxp3 molecule subject to TRAF6-mediated ubiquitination, we also screened Foxp3 constructs with various K-to-R mutations for resistance to modification by this enzyme. As before prevalent ubiquitation was seen in the Foxp3 pool in the presence of TRAF6. As expected, we also found that mutating all twenty lysine residues (20R) largely prevented the ubiquitination of Foxp3 by TRAF6. Individual constructs encoding unique single lysine residue-containing Foxp3 variants (e.g. K8, K31, K144, etc.), could, to varying degrees, support Foxp3 ubiquitination in the presence of TRAF6 (**Fig. 4A**). Of these single-lysine mutants, however, K262 was alone able to fully support TRAF6-mediated ubiquitination of the Foxp3 target protein. In contrast, mutation of Foxp3 at K262 (K262R) ablates TRAF6-mediated ubiquitination of Foxp3 (**Fig. 4B**). These results shed light on the nature of the TRAF6-Foxp3 interaction, and in particularly identify the specific residue on the Foxp3 molecule modified as a result of this pairing.

**Fig. 4.**
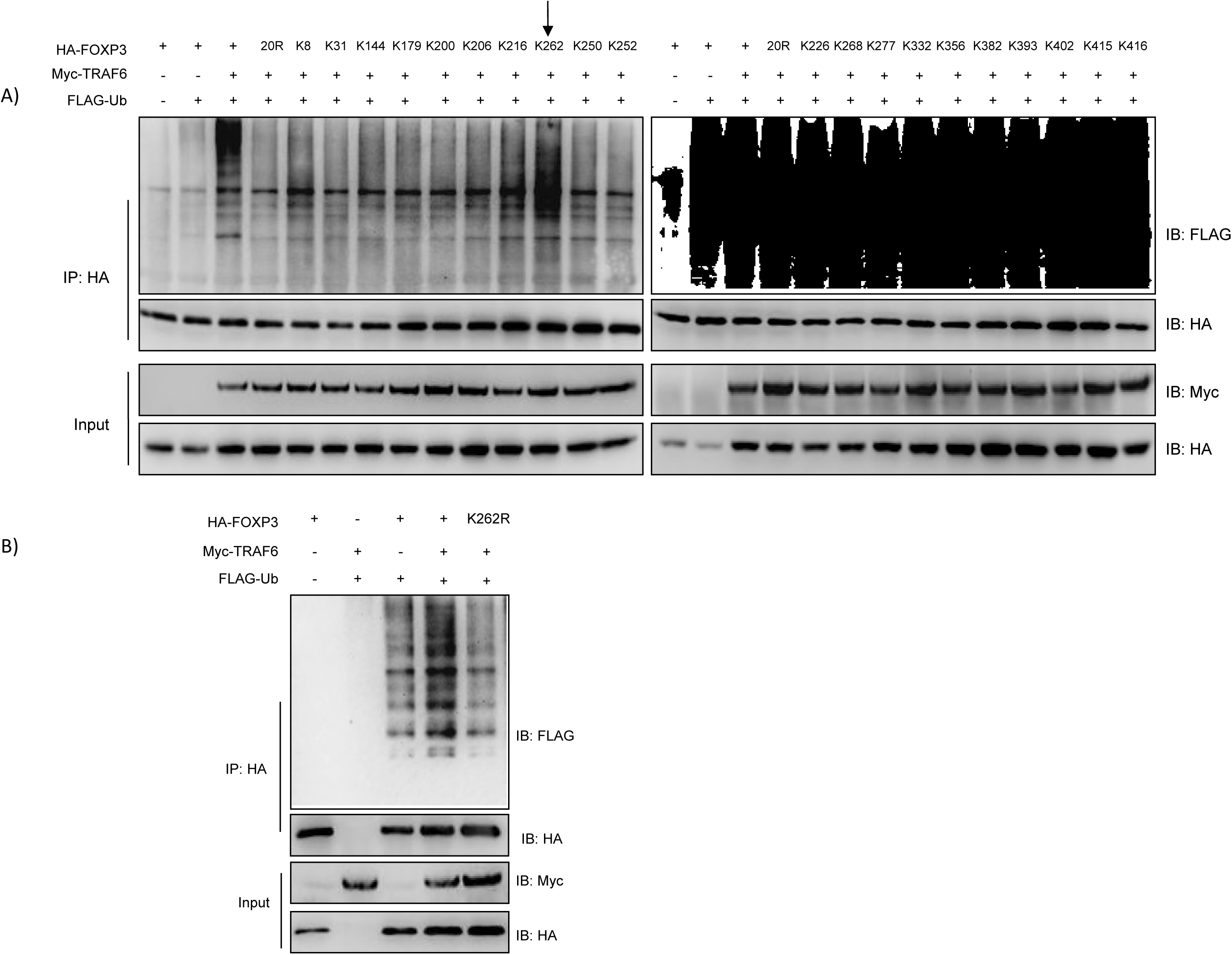
TRAF6 targets lysine residue 262 on Foxp3 for ubiquitination. (A) Immunoblot analysis of TRAF6-mediated ubiquitination of wild type, and mutant Foxp3 molecules. 293T cell lines were transfected with a normal HA-tagged Foxp3 construct, another encoding a ubiquitination resistant a mutant in which all lysine residues were replaced by arginines (20R), or one of 20 single lysine-containing construct with only the indicated lysine residue available for modification. These cell lines also received vectors for expression of TRAF6 and ubiquitin molecules labeled with Myc and Flag tags, respectively. Negative controls did not receive TRAF6 or expressed Foxp3 alone. Labeled Foxp3 proteins were pulled down from cell lysates and ubiquitinated species were visualized by immunoblotting for Flag. (B) Assessing the impact of K262 mutation on Foxp3 ubiquitination by TRAF6. As in A, 293T cells expressing combinations of Myc-TRAF6, Flag-ubiquitin, and an HA-tagged Foxp3 molecule that was either normal or possessed a K-to-R mutation at residue 262. Foxp3 proteins were pulled down with anti-HA beads and ubiquitinated proteins or total Foxp3 proteins were detected by probing for Flag and HA, respectively. Shown are representative blots from 3 experiments.

### Traf6 target residue K262 is required for nuclear Foxp3 localization and regulatory function

We then assessed how the loss of K63 ubiquitination affected the function of Foxp3. A gene reporter assay revealed, expectedly, that sequestering Foxp3 in the perinuclear region severely undermined the ability of Foxp3 to regulate gene expression. IL-2 expression is known to be among the genes directly suppressed by Foxp3 in Tregs (32-33). We therefore compared the ability of normal Foxp3 and the K262R mutant to silence expression of a luciferase reporter gene under the control of the *Il2* gene promoter. While a normal Foxp3 expression vector significantly blocked the expression of the luciferase reporter, the K63-ubiquitin-resistant K262R mutant failed to do so – resembling an empty vector (no-suppression) controls (**Fig. 5A**). This indicated that without K63 modification, Foxp3 is functionally impaired.

**Fig. 5.**
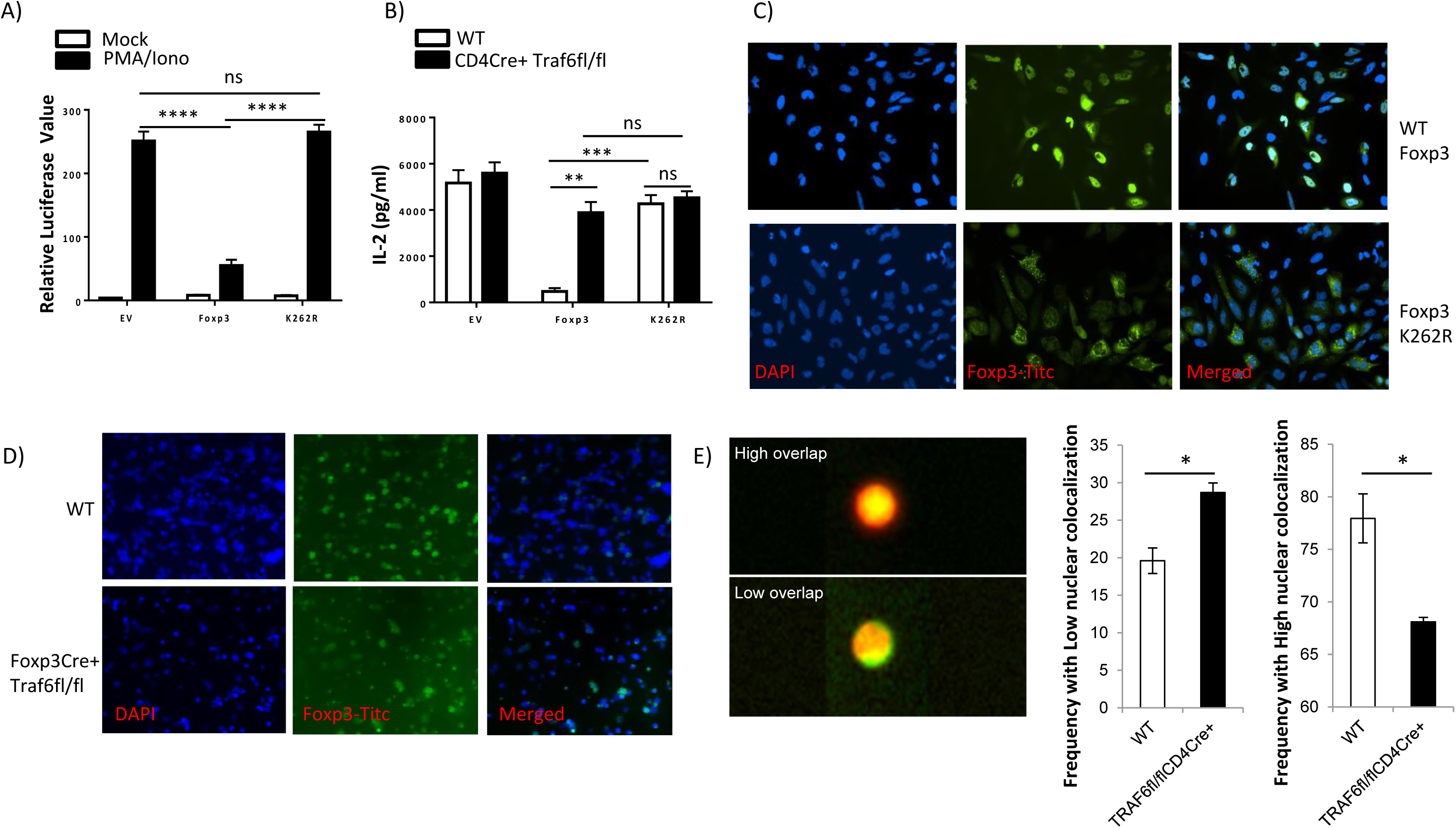
Ablation of the K63-type ubiquitination site in Foxp3 disrupts its cellular distribution, expression, and regulatory function. (A) Impact of K63-ubiquitination loss on Foxp3’s gene silencing capacity. Jurkat T cells transfected with a wild type Foxp3 expression vector, one encoding the K262R mutant, or an empty (control) vector also received a duel luciferase reporter construct where luciferase expression was under the control of the *Il2* promoter. After activation with PMA and ionomycin for 8 hours, luciferase activity was assayed. (B) IL-2 production in the presence of wild type and K63- ubiquitination resistant Foxp3. Naïve CD4+ T cells isolated from Traf6^fl/fl^CD4Cre^+^ or WT mice were activated overnight by anti-CD3/CD28 antibodies, then transduced by retroviruses carrying normal Foxp3, the K262R mutant, or an empty vector. Ires-GFP in the retroviral vector serves as an internal control. GFP+ cells were sorted out 48 hours post-transduction. The sorted cells were cultured for one additional day. Culture supernatants were collected for measuring the levels of IL-2 by ELISA. (C) Cellular distribution of wild type and K262R Foxp3. Hela cells were transfected with expression vectors encoding either wild type Foxp3 or the K262R mutant. Localization of Foxp3 protein relative to DAPI-stained nuclei was observed by fluorescence microscopy. Shown are representative (40X) fields. (D) Immunostaining of Foxp3 in Traf6^fl/fl^CD4Cre^+^ and WT derived suspensions of lymph node and spleen cells. (E) Quantification of Foxp3 distribution in murine Tregs. The degree of Foxp3-nuclear coIocalization in CD4+/Foxp3+ cells from Traf6^fl/fl^CD4Cre^+^ and WT mice was determined by Imagestream analysis, and the frequencies of cells with a low and high probability of nuclear Foxp3 distribution are shown. \**P*<0.05; \*\**P*<0.01; \*\*\**P*<0.001; \*\*\*\**P*<0.0001; ns, no significance, unpaired t test. Depicted are the results from at least three experiments. E depict mean +/-SD. A, B and F represents mean +/- SEM.

We confirmed these results in primary murine CD4+ T cells by ectopically expressing either a normal Foxp3 construct or that encoding the K262R mutant (delivered by retroviral transduction), activating the recipient cells, and measuring IL-2 production by ELISA. We found that while normal Foxp3 expression could effectively repress IL-2 transcription by wild type naïve CD4+ T cells, K262R expression could not. Importantly, when TRAF6 was absent, even delivery of wild type Foxp3 expression vector could not suppress IL-2 production (**Fig. 5B**). This nicely illustrates the importance of both TRAF6 and its target residue for proper control of gene transcription by Foxp3.

Since K63-type ubiquitination can dictate the intracellular distribution of target proteins and TRAF6 has previously been implicated in promoting K63 type ubiquitination and nuclear localization of another protein, NRIF(34), we hypothesized that modification of Foxp3 by TRAF6 also promotes the nuclear localization of Foxp3. To test this, we generated Hela cells expressing the K262R Foxp3 mutant that was resistant to K63 ubiquitination even in the presence of TRAF6. Using an immunofluorescence (IF) microscopy approach, we observed the K262R mutant was not able to localize to the nucleus unlike WT Foxp3 (**Fig. 5C**). These results suggest that TRAF6-facilitated modification at Foxp3’s K262 site is a molecular event central to the proper intracellular localization of this important transcription factor. Interestingly, when we stained Hela cells containing both the Foxp3-K262R and TRAF6 constructs, we observed that K262R mutant Foxp3 was still capable of associating with TRAF6 in the perinuclear region (**Fig. S5**) suggesting that the interaction of Foxp3-TRAF6 and the enzymatic modification of Foxp3 are indeed distinct molecular events.

Importantly, we also observed perturbations of Foxp3’s intracellular distribution in the absence of TRAF6 activity in primary murine T cells. Here, Tregs isolated from Traf6^fl/fl^Foxp3Cre^+^ mice displayed aberrant, perinuclear accumulation of Foxp3 while their normal littermates did not (**Fig. 5D, E**). Additionally, the dysfunction of TRAF6-deficient Tregs in our tumor experiments was found to be associated with this uncharacteristic distribution of Foxp3 protein. Visualization of intracellular Foxp3 protein in among the splenocytes and tumor-infiltrating leukocytes from tumor-bearing TRAF6^flox/flox^Foxp3Cre^+^ mice revealed that most of the Foxp3 detected was found in extranuclear deposits, in stark contrast to the nuclear staining expected and seen in wild type controls (**Fig. S6**). These results suggest that potential therapies targeting TRAF6-mediated K63 ubiquitination may be highly effective at breaking tolerance and bolstering the anti-tumor immunity by undermining Treg function.

### Tregs insensitive to K262 ubiquitination are dysfunctional in vitro and in vivo

These findings strongly suggest that ubiquitination of Foxp3 at K262 is key for this factor’s ability to anchor the gene expression patterns responsible for Treg function. In line with both compromised Foxp3 function and stability in the absence of K63-ubiquitination, we found that cells expressing the K262R Foxp3 mutant were broadly dysfunctional in assays of suppressive function. In vitro, CD4+ T cells transduced with lentiviral Foxp3 effectively dampened the proliferation of co-cultured naïve responder cells. In contrast, Foxp3K262R expressing cells were less effective suppressor cells (**Fig. 6A, B**). We also explored the impact of K63 resistance on Treg function in vivo using a T cell-dependent mouse model of colitis. Here, adoptive transfer of naïve CD4+ T cells (Thy1.2+) into Rag2-/- mice induced gut inflammation and progressive wasting disease in recipients. Co-injection of normal Tregs along with the colitogenic T cells effectively prevented the development of disease. CD4+ Tregs transfected to express wild type Foxp3 were also largely protective in this model. K262R Foxp3 transductants, on the other hand, failed to prevent severe colitis and mice receiving these “engineered Tregs” resembled no-Treg controls in terms of weight loss. These trends were also seen in the degree of histopathology observed across these in the large bowel (**Fig. 6C-E**). In line with these observations, mice that received no Tregs harbored considerable numbers of expanding T effector cells infiltrating their spleens, mesenteric lymph nodes and lamina propria. Meanwhile numbers of these potentially inflammatory cells (CD90.2^+^CD3^+^CD4^+^CD44^+^) were restrained in recipients of both isolated Tregs and lentiviral Foxp3 expressing cells, but largely not when expressers of the K262R mutant Foxp3 were injected (**Fig. S7A**). Also the frequencies of these potentially inflammatory cells were consistently restrained in recipients of both isolated Tregs and lentiviral Foxp3 expressing cells (**Fig. S7B and C**). Reinforcing the notion that modification at K262 is important for stable Foxp3 expression, across the tissues involved, recipients of the K262R mutant saw marked loss of Foxp3 staining in the injected (Thy1.1+) “Treg” population. In contrast, most wild type Foxp3 transductants recovered in this experiment still expressed Foxp3 (**Fig. S7D-F)**. These results further suggest that ubiquitination of Foxp3 at K262 is important for Treg function and stability. Additionally, since the overall numbers of K262R-carrying cells recovered from recipient mice were comparable to the wild type Foxp3 group (**Fig. S7D**), this modification is not likely necessary for in vivo fitness. Lastly, preventing Foxp3 ubiquitination at K262 resulted in a failure to suppress production of the proinflammatory cytokines IFNγ and IL-17 by T cells **(Fig. S7G and H)** – an observation supporting the notion that an effective and functional stable suppressor population depends heavily upon K63-type ubiquitination of Foxp3.

**Fig. 6.**
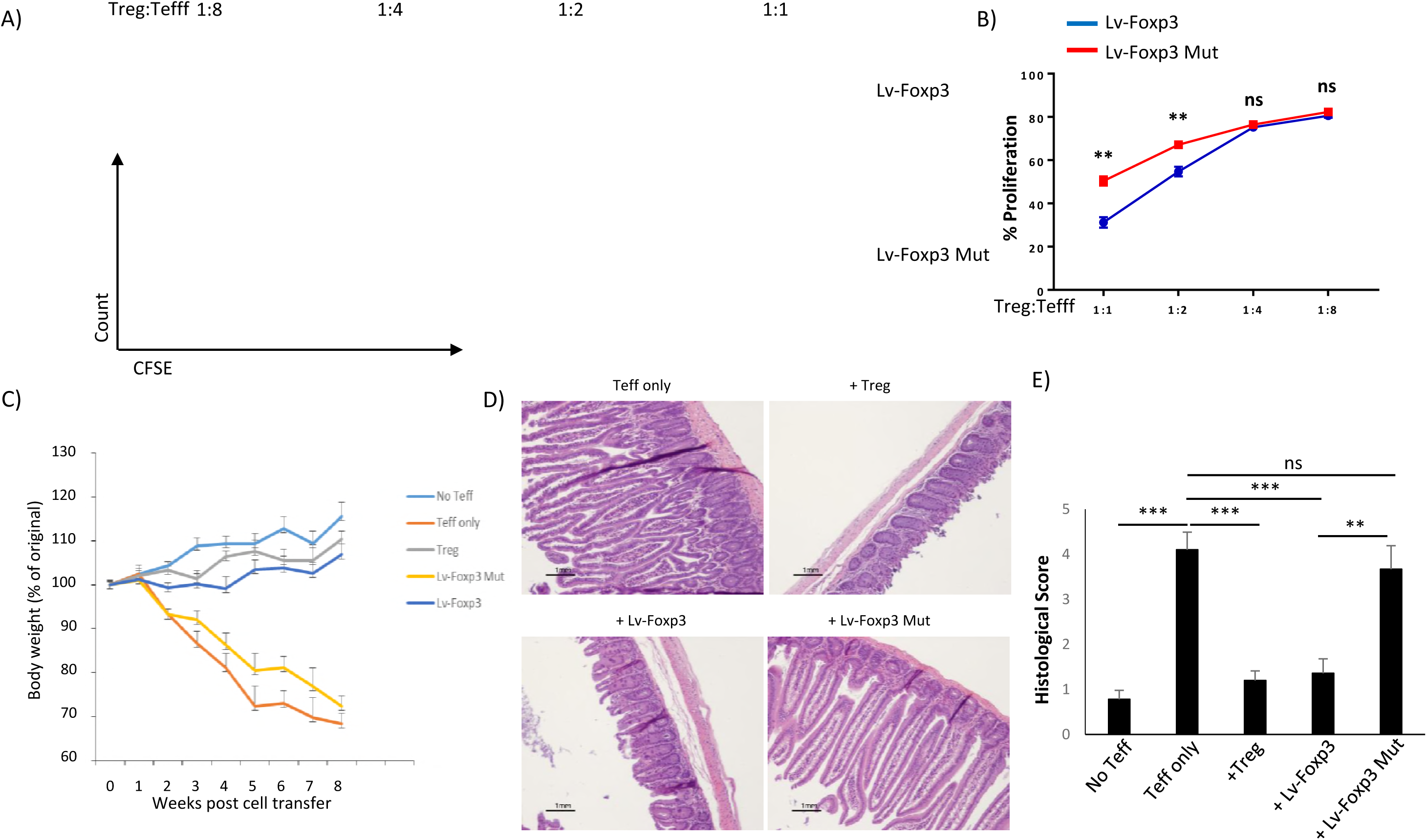
Genetic prevention of ubiquitination at Foxp3 residue K262 disrupts immune suppression *in vitro* and *in vivo*. (A, B) The in vitro suppressive potency of Wild type- and K262R Foxp3-expressing T cells. Naïve CD4+ T cells were purified from Thy1.1+ BALB/c mice and subjected to retroviral transduction to express either wild type Foxp3 or a K262R mutant resistant to ubiquitination that lysine residue 262. These engineered “Tregs” were then co-cultured with naïve responder CD4+ T cells stained with CFSE at the indicated ratio and activated with anti-CD3/CD28 antibodies. (C) *In vivo* suppression of colitis by wild type- and K262R mutant-Foxp3 expressing T cells. 2×10^5 naïve CD4+ transductants from A were mixed with 1x10^6 colitogenic naïve CD4+ T cells from wild type mice and transferred i.v. into Rag2-/- mice. Rag2-/- mice receiving no cell transfers, naïve CD4+ T cells alone, or purified wild type Tregs served as controls (n= 6/group). Recipient mouse body weights were monitored weekly for each group (D). After 8 weeks, colons were excised and processed for H and E staining (E) and histopathology scoring. **P<0.01; ***P<0.001; ns, no significance, unpaired t test. Panel A and the micrographs depicted in B are representative findings, and A (right), panels B and C represent means+/-SEM from at least two experiments.

## Discussion

Sustained Foxp3 expression is a defining characteristic of Tregs and is necessary for their function of maintaining immunological self-tolerance. Tregs are subject to a number of stabilizing or destabilizing factors via the regulation of Foxp3 fate. Previously, we and our colleagues uncovered a pathway for the degradation of Foxp3 protein hinging on K48-type polyubiquitination (15-16). This is the most important finding I believe that we characterized a distinct, non-proteolytic, pathway of K63-type ubiquitination mediated by TRAF6 that is responsible for enforcing the proper trafficking and function of Foxp3.

TRAF6 is an E3 ligase well-known for mediating K63-type ubiquitination and a widely studied member of the Tumor Necrosis Factor (TNF)-associated factor family of adaptor molecules. It participates in a number of signaling pathways triggered by the TLR/IL-1 family of receptors as well as other members of the TNF receptor family members. These pathways can be crucial for the activation of transcription factors including NFκB and AP-1 (35). Despite this association with pathways critical to immune activation, many studies have revealed a role for the E3 ligase in immune control. For instance, TRAF6-deficient mice are prone to autoimmunity (21, 36) suggesting a key role in the regulation of immune activation.

TRAF6 has also been specifically described as important for Tregs. Genetic deletion of this ligase reduces the frequency of Foxp3-expressing single positive CD4+ in the thymus (23).

Since the molecules involved in TCR-triggered NFkB activation (i.e. CARMA1, c-Rel, etc.) have been shown to be necessary for thymic Treg development (37-39), a positive role for TRAF6 in the initial upregulation of Foxp3 was proposed. Moreover, Treg frequencies in the peripherial lymphoid tissues of global TRAF6 knockout mice were also reported to be reduced compared to wild type mice, while, curiously, in vitro induction of Foxp3 expression in T cells by TGFβ was more robust in the absence of TRAF6 expression (37). While these prior findings suggested a role for TRAF6 potentially centered around the development of Tregs in the thymus, an additional role for this E3 ligase was recently identified – ensuring the phenotypic stability of Tregs.

Recently Muto and colleagues reported on the consequences of Treg-specific TRAF6 deficiency. In line with our own results they found Traf6^fl/fl^Foxp3Cre^+^ mice to be prone to dermatitis, lymphoproliferation and wide-spread immune pathologies (22). Also, they found that Tregs from these mice were only modestly less suppressive in vitro than their wild type-derived counterparts, but phenotypically unstable and ineffectual suppressors in vivo (22, 37). In accord with these observations we also found that Treg-specific ablation of TRAF6 resulted in profound in vivo defects. Interestingly, Muto et al. found that Traf6^fl/fl^Foxp3Cre^+^ mice harbored greater proportions and numbers of Foxp3-expressing T cells than wild type mice in several tissues at baseline (22). It is possible, especially given the unstable expression of Foxp3 convincingly shown by these authors that, with time, enhanced activation and exposure to inflammatory cues, Foxp3 expression in the TRAF6-deficient Treg pool may suffer. Thus differences in sampling time may account for this discrepancy. Nevertheless our present findings are in accord with those of Muto et al. as they resoundingly demonstrate that without TRAF6, Foxp3 protein levels and the gene expression pattern typical of Tregs are poorly maintained to the detriment of immune control at baseline and in distinct disease models (i.e. colitis and cancer).

Beyond confirming the importance of TRAF6 as a stabilizer of the Treg phenotype and adding to the list of scenarios wherein TRAF6 is critical for the enforcement of tolerance, the present study identifies the mechanism underlying this molecule’s role in Tregs. Specifically, we bring to light that this E3 ligase facilitates a previously unknown mode of posttranslational control over Foxp3 expression and function. We furthermore present a characterization of this K63-type ubiquitination-dependent mechanism contributing to the proper cellular distribution of this central molecular player in Treg gene expression and function.

We recently suggested positive relationship between TRAF6 expression and Treg function. We found that by antagonizing a specific microRNA (miR-146b), which targets TRAF6 transcript, they could enhance Foxp3 protein levels. Moreover, the in vitro and in vivo function of Tregs could be enhanced through a miR-146b antagomir (24). Interestingly, while increased Foxp3 expression and Treg function was attributed to an enhanced NFkB activity, which can drive transcription of the *Foxp3* gene (22, 37), levels of Foxp3 mRNA were not measured in this study. In light of our findings, it seems likely that TRAF6 contributes to both transcriptional and protein-level regulation of Foxp3 and Treg function.

Much has been elucidated regarding the transcriptional control of Foxp3 expression. For instance, transcription at the *Foxp3* gene can be dictated by epigenetic mechanisms and the interaction of multiple transcription factors (including Foxp3 itself) with several regulatory conserved non-coding regions (CNS1, CNS2, CNS3), each with unique functions(8). Additionally, it now appreciated that Foxp3 function is largely facilitated by a number of co-regulator molecules such as Eos, IRF-4, GATA-3, and other members of the Foxp3 “interactome” (4-6, 40). Posttranslational modification capable of influencing Foxp3 expression and activity are just now becoming clear (27). Surprisingly, given the undisputed importance of Foxp3 and Tregs for the maintenance of immune homeostasis, relatively little is known concerning how Foxp3 protein distribution is controlled in Tregs. The present study reveals a previously unforeseen mechanism ensuring the accumulation of Foxp3 in the nucleus that hinges on the posttranslational modification of Foxp3 protein by TRAF6.

This ubiquitin-mediated control over Foxp3’s cellular distribution and activity represents yet another opportunity for novel therapies aimed at modulating immune control. Our results present a compelling case for TRAF6 as a potential therapeutic target in anti-cancer immunotherapies. Given that TRAF6 deficiency in Tregs effectively prevented the establishment of tumor-enforced immune tolerance, pharmacological targeting of TRAF6 may be a potent means to improve anti-tumor immunity. While the potential drawbacks of such an approach could include the unintended stifling of anti-tumor effector cell activation, our findings seem to dispel this concern. Deleting TRAF6 across all T cells did not adversely affect the mounting of a robust anti-tumor response – a finding in line with the notion that, TRAF6 is not necessary for effector T cell responses, but is critical for Treg-mediated immune restraint. In light of this and our other findings, the development of therapeutic TRAF6 inhibitors will likely yield highly effective agents for sabotaging this posttranslational mode of regulating Foxp3 activity to fully unleashing the anti-tumor immune response.

In all, these findings reveal TRAF6 as a previously unappreciated posttranslational mediator of Foxp3 expression and activity in Tregs. They also provide insights into the molecular details of this distinct, non-proteolytic, and functionally enhancing brand of ubiquitination by TRAF6 including the specific modification site in the Foxp3 molecule. Importantly, this newly uncovered mode of control over Treg function may represent a target for future immunotherapies aiming to overcome immune suppression in cancer patients.

## Materials and Methods

### Mice

Traf6flox/flox mice on a C57BL/6 background were obtained from Dr. M Pasparakis (43). Foxp3yfpCre transgenic mice (originally generated in the lab of Dr. Alexander Rudensky) and CD4Cre transgenic were obtained from the Jackson Laboratory. These mice were crossed to generate mice specifically lacking TRAF6 in their Tregs. Thy1.1+ and Thy1.2+ mice on a BALB/c background and CD45.1+ C57Bl/6 mice were purchased from Jackson laboratory. All mice were housed in a specific pathogen free facility in accordance with institutional guidelines.

### Generation of mutant Foxp3 constructs

Single lysine and targeted residue mutant in Foxp3 constructs were generated by Site-Directed Mutagenesis Kits (Thermo Fisher Scientific).

### Cell line transfections, Co-IP, and Immunoblotting

To ectopically express wild type of mutant constructs in 293T cells, we utilized tagged expression vectors and were transfected by lipofectamine 2000. For Co-IP experiments, cells were lysed in RIPA buffer containing 50 mM Tris/HCl, pH 7.4, 1% Nonidet P-40, 0.5% Nadeoxycholate, 150 mM NaCl, 1 mM EDTA, with 1 mM PMSF, 1 mM Na3VO4, 1 mM NaF and protease inhibitor (Sigma), followed by immunoprecipitation with the indicated antibodies, separation by SDS/PAGE, and analysis by Western blotting. Where applicable, band density indicating protein amount was quantified using Image J software.

### Retroviral transduction of primary T cells

Naïve T cells isolated from Wild type or Traf6fl/fl/CD4Cre+ mice were stimulated with plate-bound anti-CD3 (4μg ml-1) and soluble anti-CD28 (1μg ml-1) with 60U ml-1 human recombinant IL-2 for 16 h. Activated T cells were transduced with viral supernatants (carring empaty vector, Wt Foxp3 or Foxp3 mutant) supplemented with 60U ml-1 IL-2 and 8 μg ml-1 polybrene, followed by centrifugation for 1h at 2,500 rpm. Cells were cultured at 37°C with 5% CO2 for an additional 48h and sorted out for ELISA analysis.

### T cell isolation and Treg suppression assays

Naïve CD4+ T cells from C57BL/6 mice were isolated from pooled lymph nodes and spleens by FACS as were congenically distinct Tregs (CD4+/CD25+) isolated from either wild type mice or the indicated conditional knockout strain prior to combination at the indicated ratios. Naïve T cells were stained with carboxyfluorescein diacetate succinimidyl ester (CFSE) and cocultured with equal numbers (2×10^5^) antigen presenting cells (obtained from the CD4-depleted fraction) in the presence of activating anti-CD3 antibody (0.5ug/ml). Tregs were added at the indicated ratios. 72 hours post-stimulation, division of naïve T cells was assesses by dilution of the CFSE signal. For in vivo suppression assay, CD4^+^CD25^−^CD62^hi^ Naïve T cells were sorted from CD45.1^+^ mice and CD4^+^Yfp^+^ Tregs isolated from Foxp3-Yfp^+^cre mice and Traf6 fl/fl Foxp3-Yfp^+^cre mice respectively. CD4^+^CD25^−^CD62^hi^ (1×10^6^/mice) Naïve T cells and CD4^+^Yfp^+^ (2×10^5^/mice) Tregs from Foxp3-Yfp^+^cre mice or Traf6 fl/fl Foxp3-Yfp^+^cre mice were co-injected via the tail vein (i.v) into B6 Rag2^−/-^ immunodeficient recipients (5:1). Seven days later euthanize mice, dissect spleens and place into separate labeled tubes of completed media as mentioned. Leukocytes recovered from recipient spleen were isolated and then stained for CD4 and CD45.1 after counting cell number.

### In vitro Thelper subset differentiation

Suspensions of murine leukocytes were obtained from lymph nodes and spleens. Naïve CD4+ T cells were then obtained by FACS based on their CD4+/CD62L^high^/CD25-surface marker profile. Cells were activated with stimulatory antibodies against CD3 and CD28 (1μg/ml and 2 μg/ml, respectively) for the generation of Th0 cells. Th1 cells were activated in the presence of IL-12 (20ng/ml) and anti-IL-4 neutralizing antibodies (10ug/ml). Th17 skewing was driven by inclusion of IL-6 (20ng/ml) and anti-IL-4 and anti-IL-12 (10ug/ml each) in activation media. iTregs were generated by activating naïve precursors in the presence of IL-2 (100u/ml) and the indicated concentrations of TGFβ.

### QRT-PCR and ELISA

RNA was isolated by a miniRNA extraction kit (QIAGEN). The cDNA archival kit (Applied Biosystem) was used per the manufacturer’s instruction. Triplicate reactions were run using an ABI Prism 7500. mRNA levels were determined by comparative CT method and normalized to GAPDH expression. IL-2 production from the sorted T cell culture supernatants was measured by IL-2 Elisa kit.

### Immunofluorescence Microscopy

Cells were spun down on to microscope slides at 1800 rpm for 3 minutes at room temperature (RT). The slides were then fixed with 4% paraformaldehyde for 15 minutes at RT, then washed 3 times with PBS, incubated with 0.5% Triton-X100 for 5 minutes before washing again with PBS. The cells were then blocked with 1% BSA for 1 hour at RT. FITC-labeled anti-mouse Foxp3 antibodies were diluted 1:60 with PBS and incubated with the cells for 1 hour at RT. After washing the slides, DAPI was used as a nuclear counterstain. Images were obtained using an Eclipse E800 microscope equipped with a DS-Qi1Mc camera (Nikon) and NIS-Element AR 3.0 software.

### B16 melanoma experiments

The murine melanoma cell line (B16) was maintained in vitro. For tumor challenge experiments, cells were trypsinized, washed and resuspended in PBS and 1×10^5^ B16 tumor cells were injected subcutaneously into the shaved flank or footpads of female Foxp3Cre+/Traf6flox/flox mice, CD4Cre+/Traf6flox/flox mice or their respective wild type littermates. Tumor volume was measured every other day using a digital caliper. At indicated time points post-injection, cohorts of mice were euthanized, and the leukocytes infiltrating the tumor, tumor-draining and peripheral lymph nodes and spleen were harvested and characterized by surface and intracellular immunostaining. Tumor-infiltrating leukocytes were recovered by percoll gradient centrifugation.

### Luciferase-based Transcription Activity and Repression Assays

The IL-2 luciferase reporter plasmid was co-transfected with a β-gal or renilla luciferase encoding plasmid into 293T cells or Jurkat T cells. The cells were lysed and analyzed using a luciferase assay, and results were normalized to renilla luciferase activity according to the manufacturer’s protocol (Promega). Results presented are the mean of three separate experiments, and the error bars indicate standard deviations from the mean.

### Flow cytometry & Imagestream analysis

For extracellular staining, harvested cells were washed and incubated in PBS containing 1% FBS containing fluorochrome-conjugated antibodies in a U-bottom 96- well plate. For intracellular Foxp3 staining cells were fixed and permeablized using ebioscience’s Foxp3 staining kit prior to staining. For intracellular cytokine staining, harvested cells were re-stimulated in PMA and Ionomycin in the presence of Golgi-Plug (BD). After 5 hour incubation, the cells were fixed/permeabilized (eBioscience) and incubated with antibodies against IFNγ, IL-17, or TNFα. For cellular proliferation, Cell Trace CFSE cell proliferation kit (Invitrogen) was used per manufacturer’s manual. Nuclear-Foxp3 colocalization in the Tregs of Traf6^fl/fl^CD4Cre^+^ and WT mice was assessed using Imagestream analysis (Amnis). Untouched CD4+ T cells were enriched from the lymph nodes and spleens of each group by magnetic bead isolation (Life Technologies), and intracellular Foxp3 and nuclei were stained with FITC-labelled antibody (ebioscience) and DAPI, respectively. Using IDEAS Imagestream analysis software, nuclear and perinuclear masks were generated and the frequencies of Foxp3+ cells in each group falling into bins designated high and low probability for nuclear localization were determined.

### Adoptive transfer-induced colitis model

Naïve CD4+ T cells from C57BL/6 mice were isolated from pooled lymph nodes and spleens by FACS and resuspended in PBS. 1×10^6 naïve T cells were injected i.v. into lymphopenic Rag2-/- mice. 2x10^5 congenically distinct, freshly isolated Tregs or CD4+ T cells transduced with wild type or mutant Foxp3 constructs were co-injected as indicated. Changes in body weight were assessed weekly, and upon conclusion of the experiment, colons were removed and fixed in 10% formalin. Five-micrometer paraffin-embedded sections were cut and stained with haematoxylin and eosin (H&E). The pathology of colon tissue was scored in a blinded fashion, on a scale of 0-5 where a grade of 0 was given when there were no changes observed. Changes associated with other grades were as follows: grade 1, minimal scattered mucosal inflammatory cell infiltrates, with or without minimal epithelial hyperplasia; grade 2, mild scattered to diffuse inflammatory cell infiltrates, sometimes extending into the submusoca and associated with erosions, with mild to moderate epithelial hyperplasia and mild to moderate mucin depletion from goblet cells; grade 3, moderate inflammatory cell infiltrates that were sometimes transmural, with moderate to severe epithelial hyperplasia and mucin depletion; grade 4, marked inflammatory cell infiltrates that were often transmural and associated with crypt abscesses and occasional ulceration, with marked epithelial hyperplasia, mucin depletion; and grade 5, marked transmural inflammation with severe ulceration and loss of intestinal glands. Transferred naïve and Treg cell populations were characterized by surface marker and intracellular staining for cytokines and Foxp3. Leukocytes recovered from recipient lymph node, spleen and lamina propria were re-stimulated before surface staining for CD4, CD45.1 and CD45.2.

### Statistics

The significance of differences was determined by an unpaired student’s t test, One-way ANOVA and Two-way ANOVA using Prism software (GraphPad). Differences were considered significant when P values were <0.05.

## Acknowledgements

We thank H. K. Lin (Wake Forest University School of Medicine, USA) for valuable reagents. F.P.’s research was supported by grants from the Bloomberg-Kimmel Institute, the Melanoma Research Alliance, the National Institutes of Health (RO1AI099300 and RO1AI089830), Department of Defense (PC130767), ‘‘Kelly’s Dream’’ Foundation, the Janey Fund, and the Seraph Foundation, and gifts from Bill and Betty Topecer and Dorothy Needle. F.P. is a Stewart Trust Scholar; J.B.’s research is supported by a grant from the Roswell Park Alliance Foundation and NCI grant P30CA016056. The L.L.’s research is supported by the National Natural Science Fund of China (grants 81571564, grant 91442117, grant 81521004 and 81522020), the 863 Young Scientists Special Fund (grant SS2015AA020932).

## Author contributions

J.B., F.P., and L.L. designed research; X.N., J.T., J.G., B.V.P., Z.C., S.N., and J.B. performed research; X.N., F.P., and L.L. analyzed data; H.S., X.W., B.L., S.Z. gave suggestion. BRB reversed the paper. X.N., J.B. and F.P. wrote the paper.

## Supplementary Fig. Legends

**SFig. S1. Proinflammatory cytokine production by T cells from mice with TRAF6-deficient Tregs.** (A, B) Suspensions of leukocytes from the spleens, peripheral lymph nodes (pLN) and mesenteric lymph nodes (mLN) were obtained from the tissues of WT and Traf6^fl/fl^Foxp3Cre^+^ mice. Cells were stimulated ex vivo with PMA and ionomycin in the presence of Golgistop for 5 hours. Surface markers CD4 and CD8 were stained prior to fixation, permeabilization (BD), and intracellular cytokine staining. Frequencies of the producers of the indicated cytokines within CD4+ and CD8+ T cells were found by flow cytometry (5 mice/group). \**P*<0.05; \*\**P*<0.01; \*\*\**P*<0.001; \*\*\*\**P*<0.0001; ns, no significance, unpaired t test. C and D represents mean +/- SEM. Shown are the representative findings from at least three independent experiments.

**SFig. S2. Treg-specific TRAF6 deficiency exacerbates the disruption of Foxp3 expression by iTreg exposed to inflammatory cues.** Naïve CD4+ T cells were FACS purified from the lymph nodes and spleens of Traf6^fl/fl^Foxp3Cre^+^ mice and WT mice and activated *in vitro* in the presence of Foxp3-inducing cytokines (TGFβ and IL-2) for 72 hrs. (A, B) Expression of Foxp3 under these iTreg-skewing conditions and upon addition of IL-6 (20ng/ml) to the media was determined by intracellular staining followed by flow cytometry. (C, D and E) After deriving iTregs in vitro from naïve precursors as previously described, iTregs were incubated with the indicated cytokines or LPS (100ng/ml) for 12 or 24 hours. Expression of Foxp3 was measured by intracellular staining followed by flow cytometry. A, C, and E (left) depict representative results from at least 3 experiments. \**P*<0.05; \*\**P*<0.01; \*\*\**P*<0.001; ns, no significance, unpaired t test. A (right), B (right) and C (right) shows mean +/-SEM.

**SFig. S3. Treg-specific Traf6 deficiency does not markedly alter *in vitro* Treg function but alters expression of some Treg-associated factors expression and reduces K63 ubiquitination of Foxp3.** (A) *In vitro* suppressive potency of Tregs isolated from WT and Traf6^fl/fl^Foxp3Cre^+^ mice. Tregs from the indicated mice were isolated by FACS as were naïve CD4+ responder T cells. Responder cells were stained with CFSE and co-cultured with Tregs at varying ratios in the presence of anti-CD3/CD28 antibodies. The extent of responder cell proliferation (dilution of CFSE signal) was assessed by flow cytometry. (B) Expression of Treg-associated factors including GITR, CD25, as well as surface and intracellular CTLA-4 by Foxp3+/CD4+ cells of Traf6^fl/fl^Foxp3Cre^+^ mice Tregs and their WT littermates was measured by flow cytometry. CD44 and CD62L, makers of activation and resting states, respectively, were also assessed. Shown are representative findings from at least three experiments. ns, no significance, unpaired t test. A (down) show mean +/-SEM.

**SFig. S4. The zinc finger and leucine zipper domains of Foxp3 are necessary for TRAF6 interaction.** Deletion mutants constructs of the gene for HA-labeled Foxp3 were generated according to the indicated schema (A), validated (B) and were used to map the Foxp3 regions critical for TRAF6 interaction (C, D). 293T cells were transfected to express these individual variants and immunoblotting of cell lysates for HA confirmed the predicted sizes of Foxp3 proteins lacking specified domains (B). Each deletion mutant as well as a full-length construct and an empty vector control was then co-expressed with Flag-tagged TRAF6 molecules. Pull-down with anti-Flag beads and subsequent immunoblotting for HA revealed which Foxp3 variants could interact with TRAF6. Shown are representative blots from three experiments.

**SFig. S5. Mutation of K262 does not prevent association with TRAF6.** Hela cells were co-transfected with expression constructs encoding Flag-TRAF6 and HA-K262R Foxp3. Cells were processed as described in *Methods* and stained with Alexa fluor 488-labeled anti-HA to visualize K262R Foxp3 and Cy5-labeled anti-Flag to detect TRAF6. Colocalization of these proteins was visualized by fluorescence microscopy and DAPI was used to mark nuclei. Micrographs shown are representative of at least 3 independent experiments.

**SFig. S6. Perinuclear accumulation of Foxp3 occurs in tumor-bearing mice lacking Treg-specific TRAF6 expression.** 1×10^5^ B16 melanoma cells were injected s.c. into the shaved flanks of WT and Traf6^fl/fl^Foxp3Cre^+^ mice. Approximately 21 days later, splenocytes were harvested, Foxp3 protein was detected by immunostaining, cells were affixed to microscope slides by Cytospin, and distribution relative to nuclei (DAPI+) was observed by immunofluorescence microscopy. Shown are representative 40X fields from at least 3 independent experiments.

**SFig. S7. Absence of K262 ubiquitination impairs Foxp3 expression, and suppressive function, but not cellular fitness in vivo.** Naïve CD4+ T cells were purified from Thy1.1+ BALB/c mice and subjected to retroviral transduction to express either wild type Foxp3 or a K262R mutant resistant to ubiquitination that lysine residue 226. These cells were co-injected into Rag2-/- mice along with Thy1.2+ naïve CD4+ T cells as in Fig. 6. 8 weeks later, the spleen-, mesenteric lymph node- and lamina propria-infiltrating leukocytes were recovered and characterized by flow cytometry. (A) The number of Thy1.2+ T effectors that were recovered from the indicated tissues of recipient mice. (B and C) The frequencies of CD90.2+CD3+CD4+CD44+ T effectors were assessed. (D, E and F) Similarly, the number and frequencies of Foxp3 expressing cells within the transferred “Treg” population (Thy1.1+) were assessed. (G and H) Proinflammatory cytokines IFNγ and IL-17 production by CD4+ lamina propria lymphocytes was analysed by ELISA. For A, B, D, E, Gand H, shown are the mean numbers of cells (+/-SEM) recovered from 6 mice per group from three independent experiments. \**P*<0.05; \*\**P*<0.01; \*\*\*\**P*<0.0001; ns, no significance, two-way ANOVA.

